# The Host Interactome of Spike Expands the Tropism of SARS-CoV-2

**DOI:** 10.1101/2021.02.16.431318

**Authors:** Casimir Bamberger, Sandra Pankow, Salvador Martínez-Bartolomé, Jolene Diedrich, Robin Park, John Yates

## Abstract

The SARS-CoV-2 virus causes severe acute respiratory syndrome (COVID-19) and has rapidly created a global pandemic. Patients that survive may face a slow recovery with long lasting side effects that can afflict different organs. SARS-CoV-2 primarily infects epithelial airway cells that express the host entry receptor Angiotensin Converting Enzyme 2 (ACE2) which binds to spike protein trimers on the surface of SARS-CoV-2 virions. However, SARS-CoV-2 can spread to other tissues even though they are negative for ACE2. To gain insight into the molecular constituents that might influence SARS-CoV-2 tropism, we determined which additional host factors engage with the viral spike protein in disease-relevant human bronchial epithelial cells (16HBEo^−^). We found that spike recruited the extracellular proteins laminin and thrombospondin and was retained in the endoplasmatic reticulum (ER) by the proteins DJB11 and FBX2 which support re-folding or degradation of nascent proteins in the ER. Because emerging mutations of the spike protein potentially impact the virus tropism, we compared the interactome of D614 spike with that of the rapidly spreading G614 mutated spike. More D614 than G614 spike associated with the proteins UGGT1, calnexin, HSP7A and GRP78/BiP which ensure glycosylation and folding of proteins in the ER. In contrast to G614 spike, D614 spike was endoproteolytically cleaved, and the N-terminal S1 domain was degraded in the ER even though C-terminal ‘S2 only’ proteoforms remained present. D614 spike also bound more laminin than G614 spike, which suggested that extracellular laminins may function as co-factor for an alternative, ‘S2 only’ dependent virus entry. Because the host interactome determines whether an infection is productive, we developed a novel proteome-based cell type set enrichment analysis (pCtSEA). With pCtSEA we determined that the host interactome of the spike protein may extend the tropism of SARS-CoV-2 beyond mucous epithelia to several different cell types, including macrophages and epithelial cells in the nephron. An ‘S2 only’ dependent, alternative infection of additional cell types with SARS-CoV-2 may impact vaccination strategies and may provide a molecular explanation for a severe or prolonged progression of disease in select COVID-19 patients.

## Introduction

The novel β-coronavirus SARS-CoV-2 emerged in 2019, spread rapidly and has created a global pandemic. While SARS-CoV-2 shows DNA sequence similarity to SARS-CoV ^1^ and to the bat β-coronavirus RaTG13, the infectivity of SARS-CoV-2 and zoonosis in humans was likely caused by an alteration in virus to host protein-protein interactions. A small number of very specific β-coronavirus to host protein interactions are required to infect human cells and complete replication. A comparison of the SARS-CoV-2 genome with the RaTG13 genome shows that only a few changes in the amino acid sequence of the virus entry protein (the spike protein) may have enabled SARS-CoV-2 infection and transmission in humans ^1^.

The virus to host interactome determines the virus’ infectivity and thus, tropism. In SARS-CoV-2, the receptor binding domain (RBD) of the spike protein binds to the host receptor human angiotensin converting enzyme 2 (ACE2) to initiate virus entry ^2,3^. Upon binding, high levels of the endoprotease TMPRSS2 cleave the spike protein (S) at the second polybasic cleavage site into S1 and S2’ to allow for membrane fusion. Once endocytosed, S2’ mediated fusion of the virion with the endosomal membrane releases the viral RNA genome into the host cytoplasm ^4^. Both ACE2 and TMPRSS2 are expressed in cells of the upper airways (bronchia and trachea), digestive tract (esophagus, ileum, colon, gallbladder) and common bile duct ^5^ where they confer high transmissibility, which may enable SARS-CoV-2 to linger in patients with long-lasting side effects that are not fully explained ^6^. It is also possible that a substantial number of SARS-CoV-2 cell infections occur without ACE2/TMPRSS2-mediated cell entry, albeit with much lower infectivity ^4,7^. An alternate cell entry may occur when cathepsins B/L in the lysosome cleave spike priming it for membrane fusion ^8^. Since the spike protein decorates SARS-CoV-2 virions as trimers, the proteoform composition of mature spike trimers likely determines the mechanism and specificity of any alternative, ACE2/TMPRSS2 independent cell entry. However, the precise role of spike proteoforms and their host interactomes in alternative paths for cell entry is unknown.

Open reading frames (ORF) that are encoded in the viral RNA genome are translated into virus proteins which replicate and efficiently pack the viral genome into nascent viral particles and support the egress of mature virus particles ^9^. Indeed, the virus interactome is primarily derived from virus-virus rather than virus-host interactions ^10^, and interactions with host proteins might be actively avoided to prevent any impairment of virus particle formation. However, individual viral proteins target cellular proteins to ward off detection by the immune system ^11^ or to nudge the cellular translation machinery towards a preferential translation of viral RNA transcripts ^12^. Additionally, the host interactome coordinates glycosylation ^13,14^, endoproteolytic cleavage ^15^, and trimerization ^16^ of the maturing spike protein which in turn influences selectivity and efficiency of viral cell entry in the next infection cycle ^17^.

Here, we identified the virus to host interactome of select SARS-CoV-2 ORFs in human bronchial epithelial 16HBEo^−^ cells ^18^. We devised a novel proteome-based cell type set enrichment analysis (pCtSEA) from the frequently used gene set enrichment analysis (GSEA) in transcriptomics ^19–21^, and revealed an expanded tropism of SARS-CoV-2 based on the host interactome of the spike protein. Several cell types that we identified are ACE2 negative but might harbor a proteome that allows infection with SARS-CoV-2. Further, we compared the host interactome of the D614 spike protein with the host interactome of the G614 spike. G614 yields approximately 5 fold more spike protein per mature virus particle than D614 ^22^ which increases transmissibility and infectivity ^23^; however, the mutation does not influence the severity of COVID-19 symptoms. Here we show that host proteins in 16HBEo^−^ cells processed D614 less efficient than G614 spike proteoforms. The differential processing of spike revealed that host interactors bound to the spike proteoform S2 might be important for an alternative virus cell entry.

## Results

15 of the 28 open reading frames (ORFs) in SARS-CoV-2 were expressed in human bronchial epithelial cells (16HBEo^−^). Bronchial epithelial cells serve as the main entry point of the virus into the lower respiratory tract and also provide the first line of defense against viral respiratory infections ^24^. We performed immunoprecipitations of the 2xStrep tagged viral proteins (Gordon et *al*. ^25^) and controls (2xStrep-GFP and MOCK) in biological triplicates and measured each in technical duplicates, yielding a total of 166 mass spectrometry experiments (Data availability). Bait proteins with interactors were immunoprecipitated using the CoPIT ^26^ method and digested with the endoprotease trypsin. The resulting peptides were bound to C18 reversed phase resin (PepSep) and separated by high pressure liquid chromatography (Evosep). Peptide identification with a timsTOFpro mass spectrometer (Bruker) and data analysis with SAINTexpress ^27^ (Sum SpC ≥ 5, detection in ≥ 3 replicate experiments, and ≥ 5 fold above background) resulted in an interactome comprising 15 viral and 189 host proteins (Figure 1A, Extended data network 1, Supplementary data 1). 41 of 189 (22%) host proteins interacted with more than one virus protein. Unique host-virus interactions were detected for 12 of the 15 viral proteins: orf3a (2 host proteins), orf9b (5), E (1), orf7a (2), nsp2 (2), nsp7 (3), nsp10 (5), or nsp13 (3). Nsp5-C145A, orf7b, orf8, and M protein each recruited more than 10 unique host proteins. The spike protein associated with 4 unique human proteins (TR150, LAMC2, FBX2, ERH) in addition to LAMB3, CALX, and TSP1 which were also enriched in immunoprecipitations with M and orf7b. None of the human proteins that bound to nsp9 and N were unique to either viral protein.

**Figure 1.**
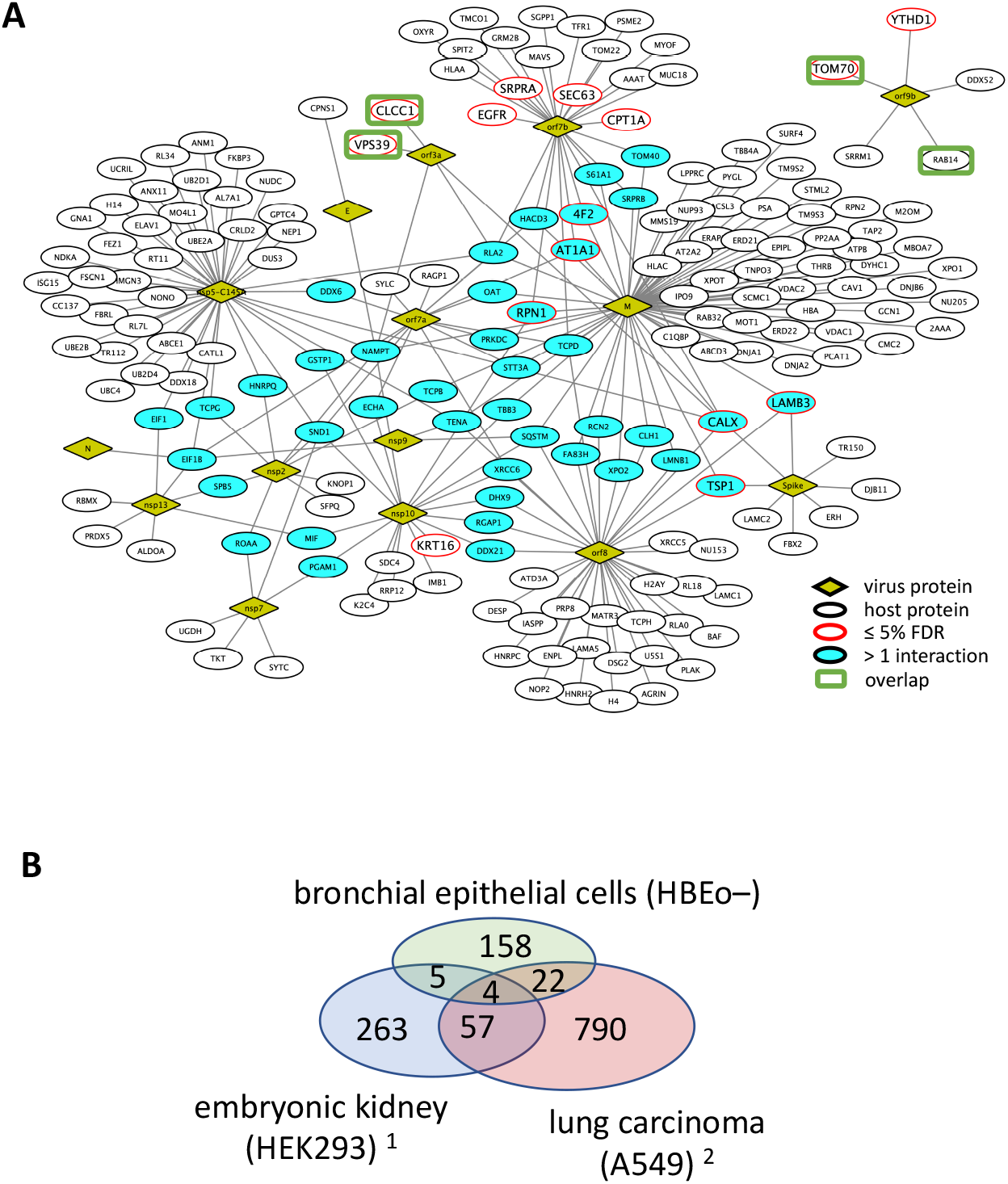
Interactome of select SARS-CoV-2 ORFs with host proteins in 16HBEo^−^ cells. **(A)** The edges in the protein-protein network indicate host proteins (ovals) that were identified in immunoprecipitations of viral proteins (diamonds). Host interactors that associated with more than one host protein or that are of high confidence (BFDR ≤ 5%) are highlighted in turquoise and red, respectively. **(B)** The Venn diagram shows the overlap of the host interactomes in HEK293 ^25^, A549 ^28^ and 16HBEo^−^ cells (this study). The green rectangles in **(A)** indicate three of the four host interactors that were identified in all three cell lines.

The overlap of the host proteins with two recently published host interactomes in human embryonic kidney HEK293 cells ^25^ and lung carcinoma A549 cells ^28^ (Figure 1B) was minimal. Common to all three interactomes were the endolysosomal trafficking proteins VSP39 (orf3a) and RAB14 (orf9b) as well as the chloride channel CLCC1, which might be involved in regulation of the unfolded protein response (UPR) ^29^, and the mitochondrial import receptor TOM70, which is targeted by orf9b to suppress type 1 interferon responses ^11^. The cell lines 16HBEo^−^ and A549 are lung airway derived, and the overlap (26 proteins, 2.5 %) was larger than between 16HBEo^−^ and HEK293 (9 proteins, 1.8 %). Therefore, we conclude that very specific virus to host protein interactions exist and that the extended virus-host interactome depends on the cellular proteome and cell type in which SARS-CoV-2 proteins were expressed. The few common SARS-CoV-2 host-virus protein-protein interactions are most likely necessary for successful virus replication.

### The SARS-CoV-2 spike interactome in 16HBEo-cells

Because the spike protein is required for successful infection, we analyzed the host interactome of spike in more detail. Two interactors, laminin-β3 (LAMB3) and laminin-γ2 (LAMC2), are extracellular glycoproteins that may impact virus cell entry or, conversely, may be functionally impaired upon SARS-CoV-2 infection. Locally confined skin blistering has been detected in conjunction with SARS-CoV-2 infection and in the presence of spike protein ^30^, and functional impairment of laminin-β3 or laminin-γ2 can cause skin blistering (epidermolysis bullosa) ^31,32^ or failure of nephrons in the kidney (Pierson syndrome) ^33,34^. Laminin-β3 can serve as co-receptor for the entry of Vaccinia virus or HPV into human cells ^35,36^.

We further found that spike binds to thrombospondin-1 (TSP1). TSP1 is present in the extracellular space and ER, and is involved in the mucosal innate immune response ^37^. TSP1 includes a C-terminal L-lectin binding domain ^38^ which may competitively inhibit a subsequent viral infection of cells as observed for HIV ^39^. Increased levels of TSP1 induce the unfolded protein response (UPR) ^40^. Key to the UPR is the ER chaperone GRP78/BiP (HSPA5), which also binds to the spike protein. GRP78/BiP might facilitate infection of epithelial cells with SARS-CoV-2 as observed for the related β-coronavirus MERS-CoV ^41^.

The extended spike interactome included DJB11. DJB11 is a co-chaperone of GRP78/BiP which binds to nascent polypeptides and stimulates GRP78/BiP dependent protein folding in the ER, or targets misfolded proteins to ERAD ^42^. Spike proteins also interacted with calnexin (CALX), which retains glycoproteins in the ER that require refolding. Calnexin was previously found to bind to the C-terminus of SARS-CoV spike protein ^43^. The F-box only protein 2, FBX2, binds N-glycosylated proteins with high-mannose oligosaccharides as part of the SKP1-CUL1-F-box E3 ligase protein complex ^44^ and targets glycoproteins to the ERAD. For example, FBX2 degrades the viral glycoprotein of the Epstein Barr virus (EBV), limiting infection of epithelial cells ^45^.

### Differences between the interactome of spike G614 to D614

After a zoonosis, viruses continue to adapt to a new host by acquiring additional mutations in the virus entry protein sequence mainly to either evade the host’s immune response or increase transmissibility. To gain insight into whether host interactors of the spike protein are altered in different SARS-CoV-2 strains, we determined the host interactome of D614G mutated spike. Mammalian expression vectors with C-terminal single or N-and C-terminal double flag tagged D614 or G614 spike cDNA were transiently transfected into 16HBEo^−^ cells and spike protein was immunoprecipitated 48 h later. Host interactors of D614 spike protein were directly compared to host interactors of G614 spike protein with CoPIT ^26,46^ (Table 1). G614 spike associated with SRP72 and high mobility group proteins (HMG). G614 but not D614 spike bound the signal recognition particle 72 (SRP72) which regulates cellular translation of membrane proteins. SRP72 relocates the translation complex with the nascent polypeptide from the cytoplasm to the rER. High mobility group proteins HMGB1 bound to G614 spike preferentially and HMGN2 or HMGB2 bound exclusively. High mobility group proteins are involved in a range of cellular pathways from proliferation to inflammation, and extracellular accumulation of HMGB1 stimulates the innate immune response to HCV ^47^.

**Table 1:**
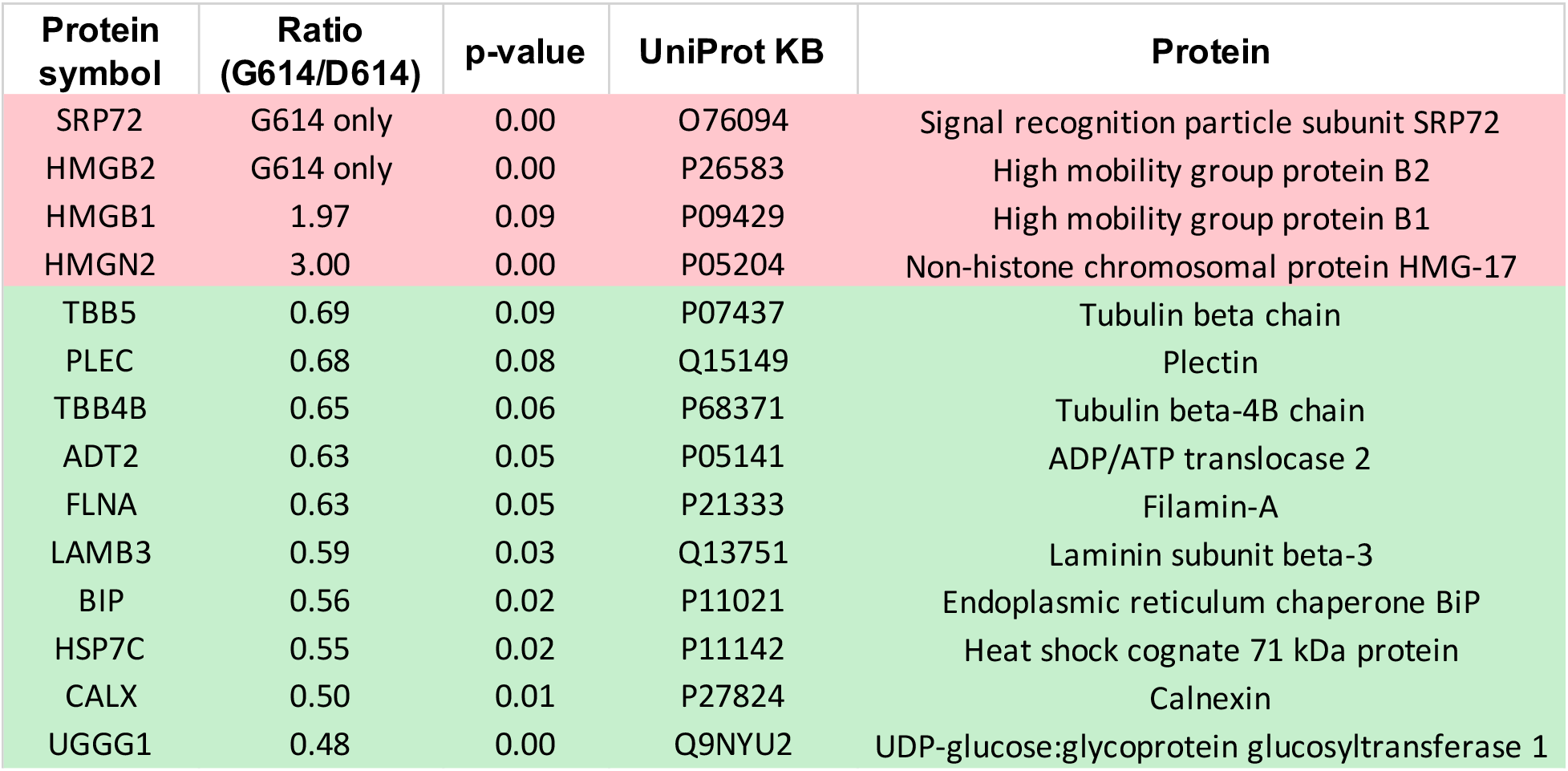
Differential binding of host interactors with spike protein variants G614 versus D614. Rows highlighted in red are host interactors that are enriched in immunoprecipitations of G614 spike whereas host interactors that preferentially bind to D614 are highlighted in green. The columns indicate the protein symbol (UniProt), the ratio of differential presence and the p-value of detection as determined with CoPIT. The UniProt identifier and name of each protein are listed separately.

D614G mutation altered the stoichiometry of the main proteoforms of spike. Spike proteoforms include S, the N-terminal S1 and the C-terminal S2 or S2’ fragment which result from endoproteolytic cleavage at the first polybasic (furin) or second (TMPRSS2) cleavage site, respectively (Figure 2A). S1 was below the level of detection in D614 spike expressing cells, whereas the mutation D614G prevented degradation of S1 and stabilized the ratio of S1: S2 at close to 1:1, as described previously ^22^ (Figure 2B). S2 was present in similar amounts to S (S2:S ~ 1:1) in both D614 and G614 spike expressing cells.

**Figure 2:**
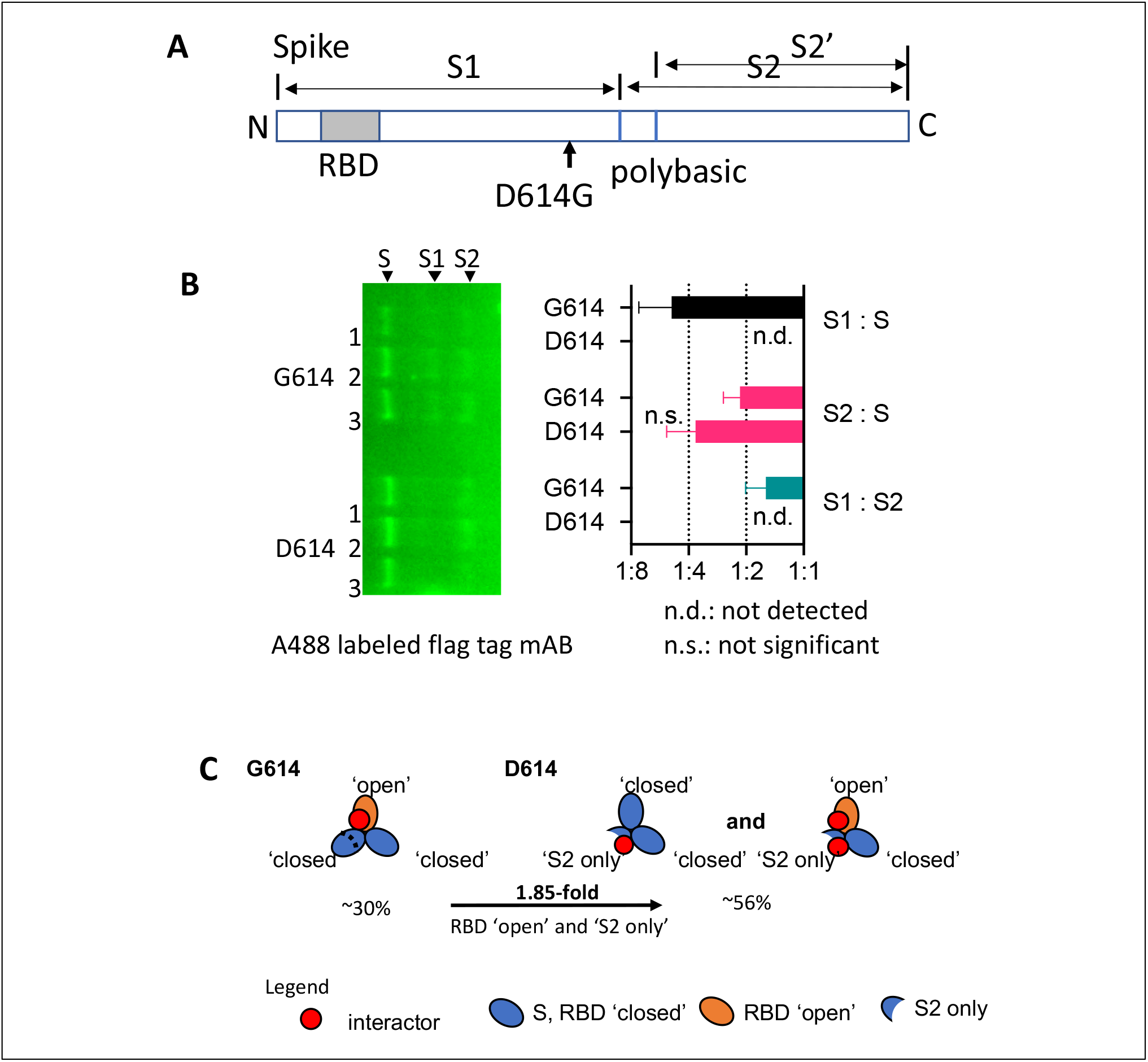
**(A)** Schematic of the spike protein and polybasic cleavage sites. Full-length spike protein (S) can be cleaved into an N-terminal proteoform S1 and two different C-terminal proteoforms S2 and S2’ which are subsumed in ‘S2 only’. **(B)** Immunodetection of flag tagged spike proteoforms with Alexa 488 labeled flag tag mAB. The bar graph indicates the relative abundance of full-length spike S and of the endoproteolytic cleavage products, the spike proteoforms S1 and S2/S2’. Full-length spike (S) harbors one anti-flag tag antibody binding site at the N-and C-terminus. **(C)** The schematic indicates the presence of host interactor binding sites on S that overlap between ‘S2 only’ and the RBD ‘open’ conformation in S. Percentages indicate the increase (~2 fold) in ‘S2 only’ binding site availability in D614 over G614 spike.

We found that heat shock cognate 71 kDa protein (HSP7C), calnexin (CALX), and UDP-glucose:glycoprotein glucosyltransferase 1 (UGGT1) were enriched by a factor of two or more in D614 over G614 spike immunoprecipitations despite the absence of S1 in D614 spike expressing 16HBEo^−^ cells (Table 1). In addition, tubulin-β (TUBB, TUBB4B), filamin-A (FLNA), plectin (PLEC), and laminin-β3 (LAMB3) were increased in D614 over G614 spike immunoprecipitations. The difference in binding can only be explained if host interactors bind to one or several cryptic motifs in S2 that are accessible only in the absence of S1 or alternatively in S with RBD in ‘open’ conformation (Figure 2C). RBD has previously been estimated to be in an ‘open’ conformation in an estimated 28% of spike proteins ^48^. Here, we used Covalent Protein Painting (CPP) ^49^ on lysine residue P0DTC2#K97 in spike, and found that RBD is in an open confirmation in ~ 20 % of G614 spike. Because D614 spike sheds S1, the number of ‘S2 only’ proteoforms is increased by approximately 33% which in turn increases the accessibility of the cryptic motifs by ~ 1.8-fold. Thus, cellular proteins that are enriched in D614 spike immunoprecipitations may bind to S2 at cryptic binding motifs that are accessible in ‘S2 only’ proteoforms that have become available by the increased shedding of S1 and in ‘open’, not ‘closed’, RBD conformations of S.

HSP7C, CALX, and UGGT1 all retain nascent proteins in the ER to ensure proper folding and refolding, or channel misfolded proteins to the ERAD. Spike D614 also bound more of the stress induced molecular chaperone GRP78/BiP. Western blots using a second set of biological replicate immunoprecipitations showed higher variability of GRP78/BiP in G614 than in D614 immunoprecipitations (Figure 3A). GRP78/BiP is exported with SARS-CoV-2 virions during virus egression ^9^ – potentially through the lysosomal pathway – and may facilitate SARS-CoV-2 cell infection as an alternative host factor that aids in virus cell entry ^50^. Recent amino acid sequence homology comparisons suggest GRP78/BiP may bind to a region in RBD in S1 ^51,52^ that encompasses the amino acid E484 in spike. Mutation (E484K) has been associated with SARS-CoV-2 mutant strains that can escape an initial humoral immune response to SARS-CoV-2 ^53^.

**Figure 3:**
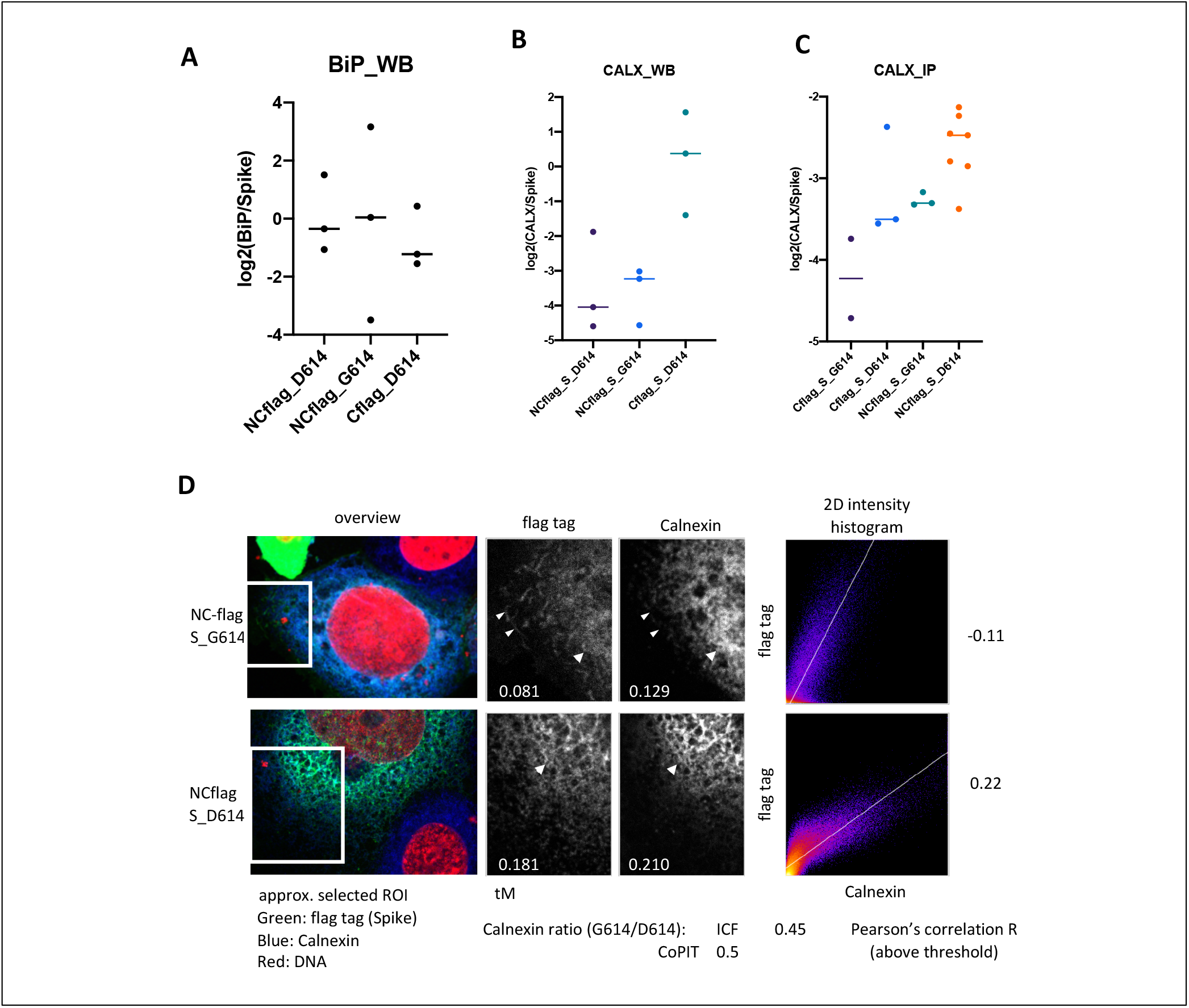
Interaction of spike with BiP and Calnexin. **(A)** The plot shows the relative abundance of GRP78/BiP to spike in three independent immunoprecipitations of spike. **(B)** Calnexin was detected and quantified in spike G614D immunoprecipitations in different biological replicate experiments by Western blot **(B)** or mass spectrometry **(C)**. Calnexin was detected by chemiluminescence (horseradish peroxidase coupled secondary antibody). Horizontal bar: Median. **(D)** Confocal images were analyzed for colocalization of spike with calnexin in immunofluorescence stained 16HBEo-cells in the regions of interest depicted in the overview. Arrows indicate colocalization of spike with calnexin whereas arrowheads point out spike protein that did not colocalize with calnexin. The colocalization of spike and calnexin above threshold was quantified with Mander’s overlap coefficient (tM) and Pearson’s correlation.

Further, we focused on the interaction of D614G spike with calnexin. Western blot analysis confirmed that less calnexin associated with G614 than D614 spike (Figure 3B), as observed with CoPIT (Figure 3C). We used immunostaining to determine the localization of spike with calnexin in 16HBEo^−^ cells (Figure 3D). A quantitative analysis of the immunofluorescence images revealed that less calnexin localized with G614 spike than with D614 spike. Moreover, G614 but not D614 spike was visible in (calnexin-negative) cellular substructures which tended to be closer to the cell membrane. Thus, maturing G614 spike might escape the ER quality control better than D614 spike that includes high levels of spike ‘S2 only’ proteoforms.

### Virus tropism based on the virus-host interactome

Tropism, defined as the cell types and tissues that can be infected by a virus, is established by the host interactome of a virus, especially in the event of zoonosis. We wanted to know whether the protein interactions of spike – and more specifically of ‘S2 only’ proteoforms in D614 spike – may lead to an ACE2 independent entry into cell types previously not associated with SARS-CoV-2 infection. To answer this question we devised a novel, proteome-based cell type set enrichment analysis (pCtSEA) to uncover SARS-CoV-2 virus tropism.

First, we sought to find out which lung cell types are most likely the target of SARS-CoV-2 based on the host proteins we identified with the 2xStrep tagged spike protein (Figure 4A). We used standard cell culture conditions to maintain proliferating cells; however, a subset constantly undergoes terminal differentiation, and ACE2 expression can be detected by RT-PCR ^54^. We determined the Pearson’s correlation of spike protein interactors with the digital gene expression (dge) in single cells isolated from human lung tissue ^55^. The original analysis included a large number of non-annotated single cells (unknown) and seven distinct lung cell types (basal, ionocytes, brush, secretory, FOXN4^+^, ciliated, and SLC16A7^+^ cells) based on differences in dge. Single cells with a correlation of *p* > 0.4 were sparsely scattered in non-annotated cells, basal and SLC16A7^+^ cells in the original single cell plot of the dataset ^55^. Overall, the dge in ciliated cells matched (*p* > 0.5) more frequently with the host interactome than with any other cell type. This initial analysis indicated that ciliated epithelial cells in the lung are likely the cell type that is most susceptible to a productive SARS-CoV-2 infection in the lung.

**Figure 4:**
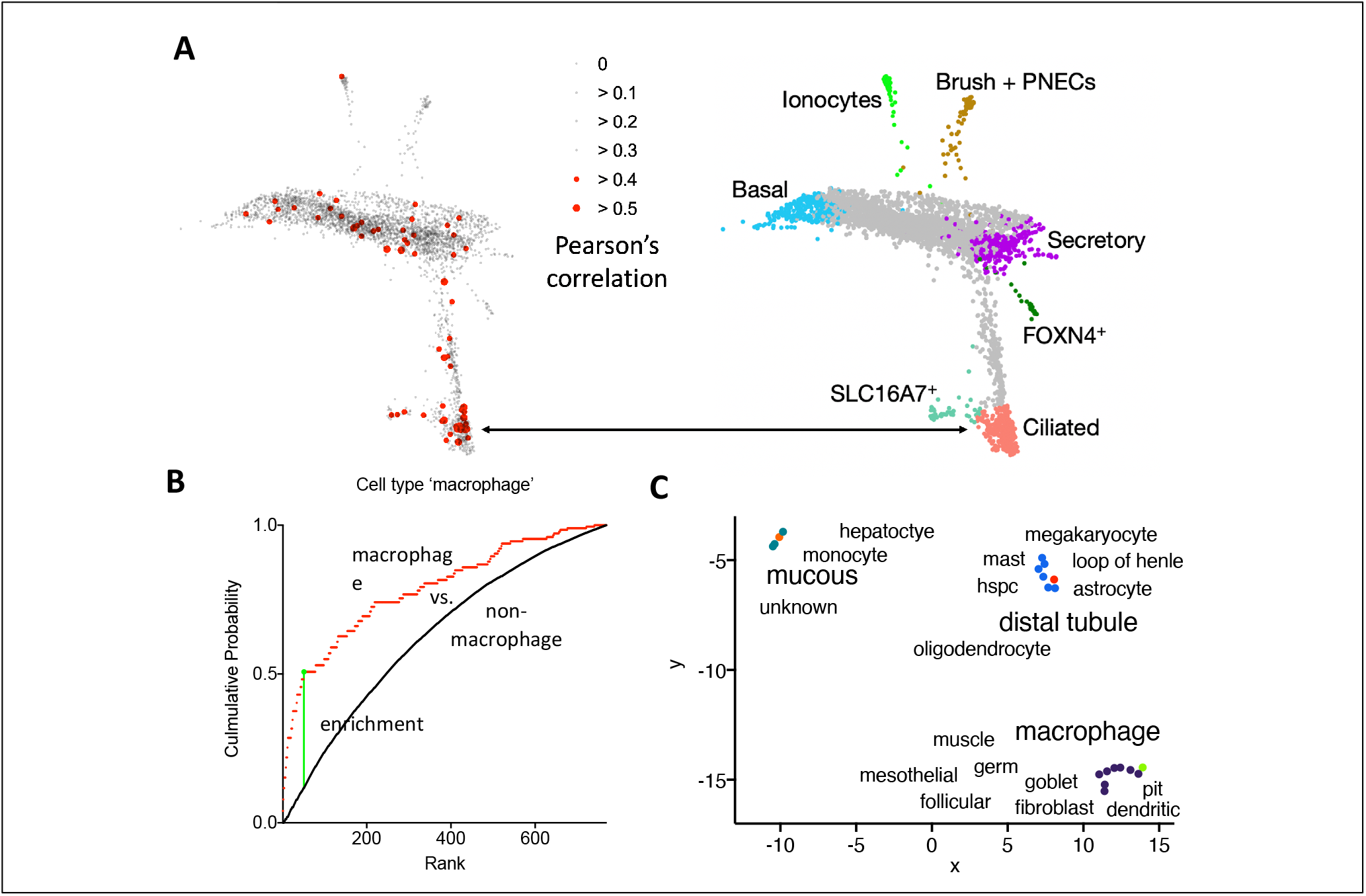
pCtSEA analysis of the SARS-CoV-2 tropism based on the spike protein interactome. **(A)** The scatter plot highlights the limited overlap of the host interactome with single cell transcriptomes of different lung cell types (adopted from ^55^). The host interactome of spike was correlated with the digital gene expression levels of single lung cell transcriptomes taking the presence and relative abundance of interactors into account. **(B)** The plot of the cumulative probability shows the weighted empirical distribution of macrophage cells in which the transcriptome correlated with the spike host cell interactome (red line), whereas the black line is the empirical distribution of all ‘non-macrophage’ cell types with *p* > 0.1. The green vertical line indicates the suprema. **(C)** Additional cell types that may be prone to SARS-CoV-2 infection are plotted in a uniform manifold approximation and projection (UMAP) which shows three clusters based on a differential presence of host interactors in the single cell transcriptomes.

Next, we improved pCtSEA so we could examine a larger panel of human single cells ^56^using a sum of host proteins identified either with flag or 2xStrep tagged spike protein. The interactome matched 918 dge patterns out of 643,636 single cell measurements with a Pearson’s correlation of *p* > 0.1. Sixty-seven of the 918 single cells were macrophages (hypergeometric test p-value < 9*10^-5^). Macrophages were significantly enriched in a two-sample Kolmogorov-Smirnov goodness of fit test of all rank ordered single cells that passed an initial Pearson’s correlation of *p* > 0.1 (K-S p-value < 0.001, Figure 4B). Seventeen additional cell ontology terms were significantly enriched (K-S p-value < 0.001) within all 117 cell ontology terms that were represented by the 918 single cells (Supplementary data 2). The remaining 99 cell types were depleted or represented by only few single cells.

The 18 cell types and cell ontology terms clustered in three different groups in a two-dimensional, uniform manifold approximation and projection map (UMAP) plot based on the single cell transcriptomes that overlapped with the host interactome of the spike protein (Figure 4C). Each group in the UMAP included one cell type that was significantly enriched when compared to a random distribution of p-values obtained by permutating the cell types in the dataset (normalized enrichment score). The significant cell ontology terms were (1) ‘distal tubule’ of the nephron or (2) ‘macrophage’ or (3) ‘mucous’ tissue-associated cells. Two of these three groups comprised cell types of similar cell ontology. For example, the group with the cell type ‘macrophage’ also included ‘dendritic cells’ which are both mononuclear phagocytes of the immune system ^57^. Cell types like ‘distal tubule’ and ‘loop of Henle’ of the nephron clustered together in the second group. Finally, monocytes and hepatocytes and cells of undefined cell ontology (‘unknown’) clustered with the cell ontology term ‘mucous’. Secretory cell types, which are characteristic of mucous tissues such as the sinus, bronchial, lung, and digestive tract epithelia, are ACE2/TMPRSS2 positive ^5^. A subset of the cell types is ACE2/TMPRSS2 negative and therefore potentially prone to SARS-CoV-2 infection in the presence of ‘S2 only’ proteoforms. Thus, an in-depth analysis of the SARS-CoV-2 spike protein interactome with pCtSEA identified three different groups of cell types that are potentially prone to a productive SARS-CoV-2 infection.

pCtSEA incorporates relative abundance of host interactors as a measure for cell type enrichment. We compared the relative expression levels of host interactors between the three groups (Figure 5). Laminin LAMB3 and LAMC2 were increased in the group labeled ‘mucous’ over ‘distal tubule’ and ‘macrophage’. A lack of filamin A (FLNA) and reduced relative levels of calnexin (CANX) but increased plectin (PLEC) characterized the group ‘distal tubule’. Thus, spike may preferentially disrupt hemi- and desmosomes formed by plectin in infected cells in the nephron. Likewise, spike might disrupt membrane anchoring of the actin cytoskeleton (FLNA) in infected cells within both groups cell types ‘macrophage’ and ‘mucous’.

**Figure 5:**
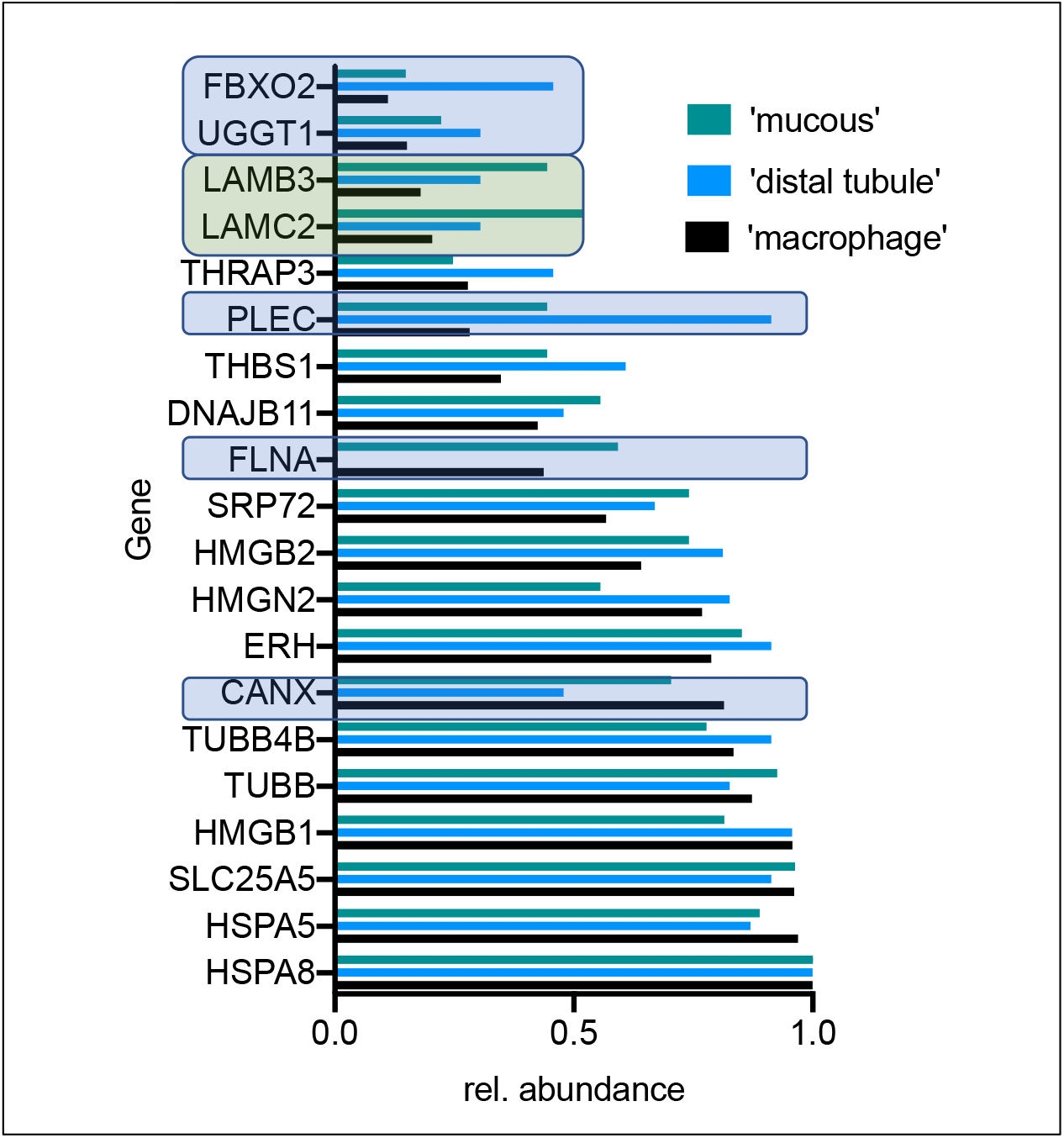
The bar graph indicates the relative abundance of gene transcripts in the three groups labeled ‘mucous’, ‘distal tubule’ and ‘macrophage’ that were identified initially with pCtSEA. Genes that deviate in expression levels relative to the group ‘macrophages’ are boxed in light blue or green for the groups ‘distal tubule’ or ‘mucous’, respectively.

## Discussion

This study revealed a small number of host proteins that specifically interact with the SARS-CoV-2 spike protein. A subset of spike protein interactors points to an ACE2/TMPRSS2 independent cell entry which might explain an infection of ACE2/TMPRSS2 negative tissues with SARS-CoV-2. We found three major proteoforms encompassing full length, non-cleaved spike protein (S) or endoproteolytically cleaved spike protein (S1-S2/S2’) or ‘S2 only’ without S1. All proteoforms are expressed in 16HBEo^−^ cells and presumably present on SARS-CoV-2 virus particles following infection of ciliated epithelial cells. Because S1 harbors the RBD, ACE2 is the preferred entry receptor during initial virus infection. However, the N-terminal domain NTD in S1 of several different coronaviruses has been shown to bind to glycosaminoglycans ^58^ or sialoglycans on glycoproteins and gangliosides turning them into co-receptors or potential restriction factors ^59^. We confirmed that D614 spike yields more ‘S2 only’ proteoforms, as observed previously. We speculate that the FBX2 associated ubiquitination complex may preferentially recognize N-glycosylated, high-mannose S1 at N234 ^14^ in D614 over G614 spike, which may target it more efficiently to ERAD. Further, we suspect that S2 in the absence of S1 may expose an otherwise cryptic receptor binding site which enables an alternative virus entry. Antibodies that bind to S2 with very high affinity and neutralize SARS-CoV-2 virus entry have been identified ^33^ and may confer protection against severe COVID-19 ^60^. Thus, we propose that loss of S1 alters the proteoform composition of spike in trimers, which in turn allows for infection of cells through an alternative, ACE2-independent virus entry. We speculate that extracellular laminins may serve as additional co-receptors to increase virus uptake.

This interpretation is supported by a pCtSEA analysis of the host interactome of spike. pCtSEA indicated that epithelial cells in the nephron and macrophages might get infected with SARS-CoV-2 in addition to lung epithelial cells. Notably, the immune cells identified with pCtSEA may be prone to antibody-dependent enhancement (ADE) of infection with β-coronaviruses ^61^ in the absence of ACE2. The cell types identified with pCtSEA reside in several different human tissues including muscle and brain (oligodendrocytes and astrocytes). We speculate that an expanded tropism of SARS-CoV-2 might shed light on a prolonged progression of COVID-19 in select patients. A pCtSEA analysis can indicate whether a virus-host interactome in its native reservoir may permit a zoonosis, as observed in the COVID-19 pandemic. Indeed, when the virus did not co-evolve with the host, a high variability of host-virus protein-protein interactions may arise from introducing viral proteins into a novel, differently equilibrated host environment. Hence, it might explain the high variability in host interactors of SARS-CoV-2 that were identified in different cell lines and tissues ^25,28,62^.

Our data analysis suggests that SARS-CoV-2 may utilize two alternate paths for cell entry: one path exploits ACE2 receptor binding, and the other path may be based on ‘S2 only’ proteoforms that may bind to laminins as coreceptors. As a consequence, an adaptive immune response that targets almost exclusively antigenic S1 may fail to completely curb severe or persistent infections with SARS-CoV-2. Therefore, the host interactome of spike suggests that immunization strategies might provide increased protection against COVID-19 if the ‘S2 only’ proteoform is also used as immunogen.

## Methods

### Expression plasmids

All SARS-CoV-2 ORF expression plasmids were obtained from the laboratory of Nevan Krogan ^63^. Viral ORFs were provided as inserts into the vector backbone pLVX-EF1alpha-2xStrep-IRES-Puro except for the spike protein cDNA which was cloned into pTwist vector and extended at the C-terminus with a 2xStrep tag. Expression vectors for the N-terminal or N- and C-terminal flag tagged spike protein variants D614 and G614 were kindly provided by Hyeryun Choe’s laboratory ^25,64^.

### Cell culture, transfection and transduction

HEK293T cells were grown at 37 °C and 5 % CO_2_ in DMEM supplemented with 10% tetracycline-free fetal bovine serum and 1% Penicillin-Streptomycin (Invitrogen). Cells were transfected with the LentiX VSV-G transfection mix (Clontech) and different SARS-CoV-2 ORFs (cloned into pLX vector). VSV-G pseudo-typed lentiviral particles were collected 48 h later and filtered through a 0.22 μm filter. Human bronchial epithelial cells (16HBEo-) were transduced with VSV-G pseudo-typed lentiviral particles in the presence of 3 μg/ml polybrene (Millipore). Expression vectors encoding the spike protein cDNA were directly transfected into 16HBEo^−^ with Lipofectamine P3000 according to manufacturer’s recommendations (Invitrogen). Transduced and transfected 16HBEo^−^ cells were grown at 37 °C and 5 % CO_2_ for additional 48 h prior to harvest.

### Immunoprecipitation

Cells were lysed in TNI lysis buffer (50 mM Tris pH 7.5, 250 mM NaCl, 1 mM EDTA, 0.5 % Igepal CA-630, 1 x Complete ULTRA EDTA-free protease inhibitor mix (Roche), 1 x HALT Phosphatase inhibitor mix (Thermo) for 20 min at 4 °C and further processed for immunoprecipitation according to ^46^. 2xStrep tagged SARS-CoV-2 ORFs were immunoprecipitated using 30 μl of Megastrep type 3 XT beads (IBA LifeSciences, Germany), and flag-tagged SARS-CoV-2 ORFs were immunoprecipitated using Anti-DYKDDDDK Magnetic Agarose (Pierce) at 4 °C for 16 h. Beads were washed 3 x with 10 volumes of TNI buffer (50 mM Tris pH 7.5, 250 mM NaCl, 0.5 % Igepal CA-630) and 2 x with 10 volumes of TN buffer (50 mM Tris pH 7.5, 250 mM NaCl, 1 mM EDTA) before elution with 8 M urea, 5 mM TCEP, 0.1 % (w:v) Rapigest (Waters) for 15 min at 60 °C.

### Digestion, sample purification and mass spectrometry

Eluted proteins were alkylated with chloroacetamide (10 mM, 30 min, 37 °C) and digested at 37 °C for 16 h in 100 mM Tris HCl, pH 7.2, in the presence of 0.5 to 0.8 M urea with the endoprotease Trypsin (Promega). Peptides were subsequently bound to the reversed phase in Evotips (Evosep) and washed according to manufacturer’s recommendations. Each immunoprecipitation was measured in technical duplicate. Following elution and chromatographic separation of peptides on a 15 cm ReproSil C18 column (3 μm, 120 Å, id 100 μm, PepSep) with a 45 min gradient (Evosep), peptides were detected and fragmented with a timsTOFpro mass spectrometer (Bruker) with PASEF enabled ^65^.

### Data analysis

Fragment ion mass spectra were extracted from timsTOFpro raw data files and stored in the ms2.txt flat-file format. Fragment ion spectra were searched with Prolucid ^66^ with the forward and reversed human UniProt database v.07-01-2020, and filtered with DTASelect ^67^ to a false discovery rate of < 1 %. Immunoprecipitation experiments were filtered with SAINTexpress to remove non-specific background proteins ^68^ (Supplementary Table 1). The final virus-host protein-protein interactome includes all 2xStrep tagged virus-host protein interactions. Due to a different background of contaminants of flag versus 2xStrep tag, flag-tag immunoprecipitations were not further considered for the spike protein in the final network of virus-host interactions immunoprecipitations (Figure 1A). Spike protein immunoprecipitations of the variants D164 and G614 were directly compared with CoPIT ^26^. Interactors present in ≥ 2 biological replicates and detected as differentially expressed with a cut off value of 0.1 (CoPITgenerator) were further analyzed. Keratins are often observed as contaminants in immunoprecipitation experiments from epithelial cell lines and therefore were excluded from further analysis.

### Immunofluorescence detection and protein co-localization

48 hours post transfection cells were washed with 1x PBS, fixed in 4% paraformaldehyde/1x PBS for 15 min at RT, permeabilized with 0.1 % Triton X-100/1xPBS for 10 min at RT, washed three times with 1x PBS and incubated with 5 % horse serum/1xPBS for 1 h at 20 °C before incubating with the following antibodies (Cell Signaling Laboratories): anti-calnexin (clone C5C9), anti-BiP (clone C50B12), anti-Hsp90a (clone D1A7), anti-ZO1 (clone D7D12), anti-ZO2 (2847), anti-ZO3 (D57G7), anti-claudin 1 (D5H1D), anti-CD2AP (2135), anti-afadin (clone D1Y3Z) or Alexa Fluor 488 anti-DYKDDDDK (flag) tag monoclonal antibody (L5, Life Technologies). The plasma membrane was stained with CF594 dye conjugated wheat germ agglutinin (Biotium), while ER was stained with the ER Cytopainter staining kit (AbCam) according to manufacturer recommendations. Images were taken with a Zeiss LSM 780 confocal laser scanning microscope and analyzed using Fiji/ImageJ (v.2.1.0/1.53c).

### Western blotting

One volume of 2x SDS sample loading buffer was added to samples in TN buffer (1:1; v:v) and samples were heated to 50°C for 15 min. Proteins were separated on NuPAGE 4-12% Bis-Tris gels (Life technologies) before transferring proteins onto Nitrocellulose membranes and incubating with anti-BiP (C50B12, Cell Signaling Laboratories (CSL)), anti-ZO1 (D7D12, CSL), anti-ZO2 (2847, CSL) or Alexa Fluor 488 conjugated anti-DYKDDDDK Tag monoclonal antibody (L5, Sigma) in 5 % BSA, 1x PBS, 0.05 % Tween-20. Signals were detected either using Radiance Q ECL reagent (Azure Biosystems) after washing and incubation with HRP coupled anti-rabbit antibodies (CSL) or imaged directly using an Azure 600 Western blot imaging system. Western blot images were quantified in Fiji/ImageJ.

### Proteome based Cell Type Set Enrichment Analysis (pCtSEA)

The input dataset for a pCtSEA analysis can be the host interactome of SARS-CoV-2 or any proteome that is listed with a UniProt identifier and an abundance value (Spectral counts, SpC). Importantly, pCtSEA assumes that at least a partial stochiometric relationship exists between the relative abundance of mRNA transcripts in cells the proteins in the interactome. First, single cell transcriptomes are filtered for the expression of ≥ 10 genes that are also present in the proteome. All protein abundances are correlated with the single cell digital gene expression (dge) values (Pearson’s correlation), and each cell type or cell ontology term with ≥ 10 cells that pass the Pearson’s correlation threshold of p > 0.1 is further evaluated. pCtSEA calculates the p-value of the hypergeometric test and an enrichment score based on a two-sample Kolmogorov-Smirnov goodness of fit test (K-S test) in which the empirical cumulative distribution function of cells specific for a cell type is compared to the remaining cells when rank ordered by their correlation values. In addition, the value and rank position of the supremum is reported. Furthermore, each cell type enrichment score is differentiated from a population of enrichment scores (suprema) that is obtained in an iterative process in which cell type labels are randomly permutated, yielding an empirical p-value that indicates the significance of enrichment. Finally, the total distribution of all enrichment scores of all cell types is corrected for multiple hypothesis testing and a false discovery rate (FDR) associated with each cell type.

Cell types that are significantly enriched (K-S test) are clustered based on the relative overlap of the proteins that correlated with the single cell gene expression pattern using a uniform manifold approximation and projection (UMAP). pCtSEA is accessible as web tool: http://pctsea.scripps.edu.

## Acknowledgements

We are indebted to Hyeryun Choe and April Tobey for the HA-tagged spike protein D614G. The thank Claire Delahunty for critically reading the manuscript.

## Author Contributions

C.B. and S.P. designed and executed the experiments. J.D. analyzed the samples on the mass spectrometer, R.P. performed data analysis of mass spectrometric data. C.B. conceived pCtSEA and S.M.-B. realized pCtSEA. J.R.Y. provided funding and instrumentation. C.B. performed data analysis, prepared figures and wrote the manuscript.

**Extended data network 1: The virus-host protein network of select SARS-CoV-2 open reding frames in 16HBEo-cells is depicted.** The bipartite network is available at the url: http://ndexbio.org/#/network/cb71d1f8-12fa-11eb-9eee-0ac135e8bacf?accesskey=1713beeb8f740c163894126d8346ea4c20edbf1e4910f4da41c6bdd153f673b5

**Supplementary data 1:** The table lists all proteins detected in the IP-MS experiments and their respective scores upon analysis with SAINTexpress for the probability of a virus-protein interaction.

**Supplementary data 2:** pCtSEA analysis of the host interactome of spike in 16HBEo^−^ cells.

## Data availability

Mass spectrometric fragmentation ion spectra and quantification files can be accessed in Massive (MSV000086433) or ProteomeExchange (PXD022457). Access will be made available upon request by reviewers.

